# Light-induced melatonin suppression in school-aged children before and during camping: associations with circadian phase and daytime light exposure

**DOI:** 10.64898/2026.11.12.766095

**Authors:** Taisuke Eto, Kota Takeoka, Kana Nishimura, Michihiro Ohashi, Sang-il Lee, Yuki Ikeda, Yuki Nishimura, Shingo Kitamura, Akiko Shikano, Shingo Noi, Shigekazu Higuchi

## Abstract

Prior light exposure can influence the magnitude of melatonin suppression in response to evening light, even within the same individual. We examined whether light-induced melatonin suppression in school-aged children differed before and during a camp with approximately two weeks of exposure to a natural light-dark cycle. We also explored interindividual variation in melatonin suppression and its associations with circadian phase, habitual sleep timing, and daytime light exposure. Twenty-one healthy children aged 9–14 years underwent evening light experiments at the start of camp (Pre-camp session) and approximately two weeks later (Camp session). Melatonin suppression was calculated by comparing the area under the melatonin concentration curve between the evening light condition (200 ± 30 lx, 5000 K) and the dim light condition (< 30 lx, incandescent lamps), with integration starting at dim light melatonin onset (DLMO). We assessed sleep habits from 14-day sleep diaries before camping and measured wrist-level illuminance during camping. We examined associations using Spearman correlations and compared Pre-camp and Camp melatonin suppression using a paired *t*-test. Melatonin suppression did not differ significantly between Pre-camp and Camp sessions (*p* = 0.21). Greater suppression in the Pre-camp session was associated with later DLMO (*rho* = 0.49, *p* = 0.028), but not with habitual sleep variables. In the Camp session, higher same-day geometric mean illuminance was associated with lower suppression (*n* = 15; *rho* = −0.58, *p* = 0.024), whereas 7-day light-exposure metrics were not. Mean melatonin suppression did not significantly change across the camping period. Meanwhile, children showed large interindividual variation in melatonin suppression, and circadian phase and recent daytime light exposure are candidate correlates of this variation in the responsiveness to the evening light.

## Introduction

Light is the principal environmental cue that synchronises the human circadian system to the 24-hour day. Light exposure in the early biological night can delay circadian phase^1^ and also acutely suppress melatonin secretion^2,3^. The modern evening environments extending light exposure after sunset and into the hours preceding sleep by artificial lighting disturb circadian rhythm and sleep timing^4,5^ and suppress melatonin secretion, even at low room-light intensity^6^. Adverse effects of late circadian rhythm and a concomitant experience of social jet lag, late sleep timing and a concomitant lack of sleep duration on human health have been well studied^7^. Light-induced melatonin suppression, together with circadian disruption and sleep loss, has been proposed as one of the potential mechanisms linking exposure to light at night with adverse health outcomes^8–10^. Therefore, it is crucial to understand the mechanisms of melatonin suppression to reduce the harmful effects of light at night.

Children’s circadian rhythm or melatonin secretion may be particularly affected by evening light^11–14^. It has been demonstrated that melatonin in school-aged children was suppressed by night light exposure more than in middle-aged adults^15^ especially in the blue-light region^16^. This greater response may be partly attributable to larger pupil size and higher crystalline lens transmittance, which increase the amount of short-wavelength light reaching the retina^17^. Although evidence of the possibility that children’s circadian system has a high sensitivity to night light has been accumulated, previous studies have primarily focused on mean differences between age groups, and interindividual variability in children’s melatonin suppression and possible factors affecting this interindividual variability have not yet been characterised.

In adults, sensitivity to evening light varies across individuals. Phillips et al. demonstrated a more than 50-fold difference between individuals in the light level required to induce 50% melatonin suppression^18^. Such variability may be related to individual differences in circadian and sleep timing. Individuals who are more sensitive to evening light may experience stronger phase-delaying effects under the same light environment and may consequently be more susceptible to delayed sleep timing. Consistent with this possibility, adult studies have reported associations between evening light sensitivity and habitual bedtime, delayed circadian phase, or later sleep timing^19–21^. Nevertheless, these relationships remain incompletely understood, and it is unclear whether comparable associations exist in children. In particular, few studies have examined whether individual differences in children’s melatonin suppression are associated with their circadian phase or with habitual sleep timing.

Melatonin suppression may also vary within an individual depending on recent light exposure. Controlled within-subject studies in adults have shown that subsequent light-induced melatonin suppression was lower after exposure to brighter light than after exposure to dim light^22,23^. In a study that assessed the same participants in summer and winter, both daily light exposure and melatonin suppression differed between seasons^24^. However, it remains unclear whether daily light exposure is associated with melatonin suppression in children under field conditions. Compared with typical indoor living, camping under a natural light-dark cycle generally involves brighter days and darker evenings. Exposure to a natural light-dark cycle has been shown to advance circadian phase in adults^25^ and children^26^. Whether the same children show different melatonin suppression responses to evening light before and during camping remains unclear.

The previously reported camping study^26^ provided a unique opportunity to examine light-induced melatonin suppression in the same school-aged children shortly before camping and approximately two weeks into the camping programme under a natural light-dark cycle. The primary aim was to determine whether mean melatonin suppression differed between the two assessments. We hypothesised that continuous exposure to the natural light environment during camping would be accompanied by a change in children’s melatonin suppression responses. We additionally conducted secondary exploratory analyses to characterise the extent and distribution of interindividual variability in melatonin suppression under daily habitual living, i.e., just before camping life, and to examine its potential correlates. Specifically, we examined whether melatonin suppression before camping life was associated with circadian phase and habitual sleep timing. We also examined whether personal light exposure measured earlier on the day of the observing melatonin suppression was associated with melatonin suppression during camping life. These exploratory analyses were intended to generate hypotheses regarding individual and environmental factors associated with children’s response to evening light.

## Methods

### Participants

Participants were 21 healthy children (18 boys and 3 girls) aged 9–14 years (mean ± SD, 10.4 ± 1.1 years) who participated in a camping program conducted from July to August 2017. None of the participants had a medical condition or was taking medication regularly. All participants provided written informed consent to participate in the study, which was approved by the Ethical Committee of Kyushu University. Informed consent forms for the children were completed by the parents after confirming their children’s agreement to participate. The experiments were conducted in accordance with the Declaration of Helsinki.

### Study design and baseline monitoring

The summer camp programs were conducted by Mobilityland Corporation in Japan. The campsite was located in Tochigi Prefecture, Japan (latitude: 36.5°N, longitude: 140.2°E). The mean sunrise and sunset times during the camping program were 4:43 and 18:49, respectively. Except on the first and second days, bedtimes and wake times were scheduled by the organiser at 20:00 and 5:00, respectively. Scheduled meal times were 6:30 for breakfast, 12:00 for lunch, and 17:00 for dinner. Outside sleep and meal periods, the children participated in camping program events, including playing in the forest, playing in a river, and craftwork workshops. During the evening hours, participants prepared and ate dinner and studied in a shed with incandescent lamps. Illuminance inside the shed in the evening was less than 30 lx.

During the 2 weeks preceding the camping program, the participants were instructed to maintain their habitual bedtime and wake time and to keep a sleep diary independently. Sleep-wake habits and daily light exposure in that 2-week period were recorded by wrist activity monitors with a light sensor to verify compliance with instructions (MotionWatch 8, CamNtech Ltd., UK). Monitoring continued during the camping program. The sleep-related and light exposure data obtained from wrist activity monitors before the camping program were excluded from further analysis because they were frequently missing: participants could not wear them at school.

Figure 1 shows an overview of the experimental procedure in this study. The evening-light experiment to observe light-induced melatonin suppression was conducted on the first and second nights of the program (Pre-camp session), before the camp schedule was expected to have modified habitual sleep timing, and on the 16^th^ and 17^th^ nights (Camp session). In the Pre-camp session, the light exposure condition (Light condition) was administered on night 1 and the dim-light control condition (Dim condition) on night 2. Participants remained in the assigned lighting from 17:00 until their habitual bedtime. The order was fixed so that the first camp evening resembled the participants’ usual indoor-light environment and to minimise the burden of an unfamiliar Dim condition immediately after arrival. In the Camp session, the Dim condition was administered on night 16, and the light exposure on night 17, thereby avoiding the circadian phase shift caused by the light exposure first. In the Camp condition, unlike in the Pre-camp condition, participants remained under the assigned lighting from 17:00 to 20:00.

**Figure 1.**
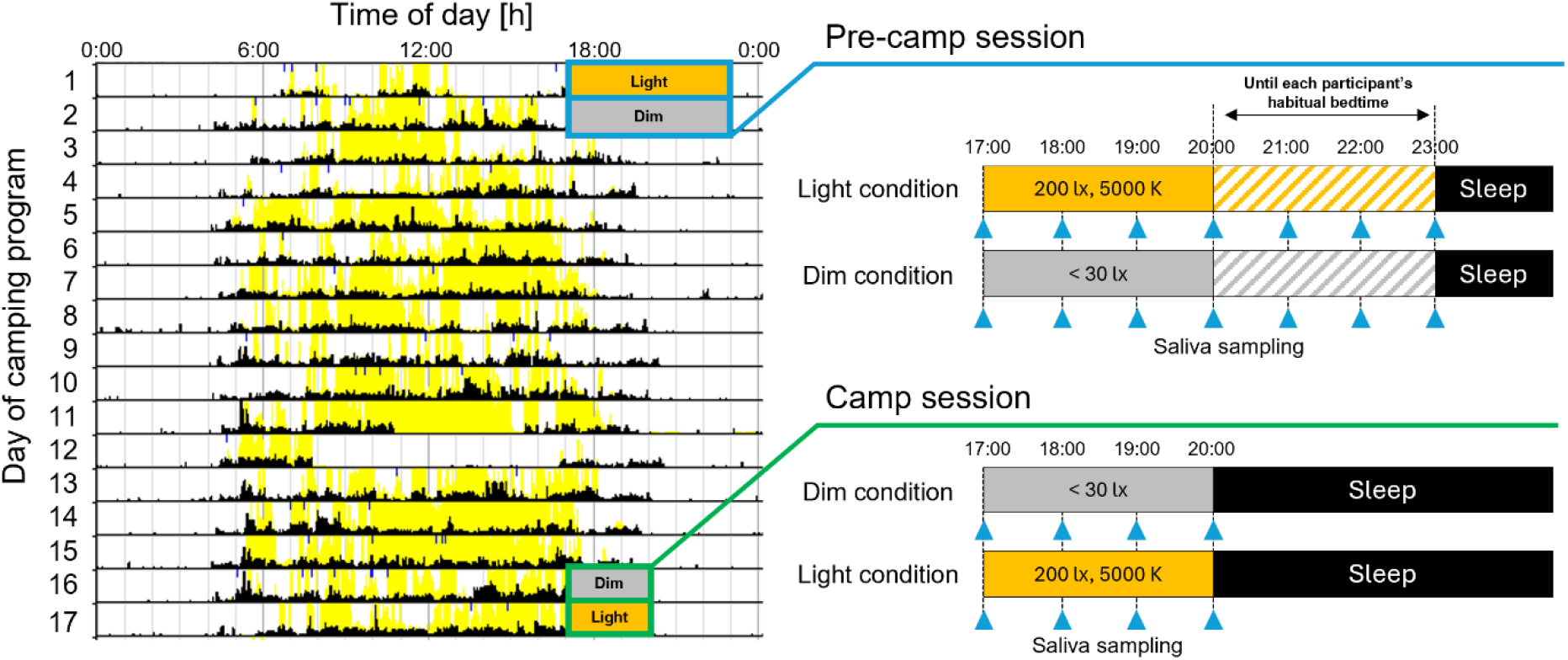
Overview of the experimental procedure. The evening-light experiment to observe light-induced melatonin suppression was conducted on the first and second nights of the program (Pre-camp session), and on the 16th and 17th nights (Camp session).

### Evening light conditions and saliva collection

In the evening-light experiment, participants were seated in a shed located at the camping site and were asked to remain near their assigned seats. Under the Light condition, six commercially available LED luminaires (HH-LC560A and HH-CB0833A, Panasonic Corp., Japan) provided a vertical illuminance at eye level when participants were seated and looking straight ahead of 200 ± 30 lx and a correlated colour temperature of 5,000 K. Under Dim conditions, incandescent lamps provided < 30 lx, measured under the same conditions as the Light condition and a correlated colour temperature of ≤ 3,000 K. All light measurements were performed using an illuminance spectroradiometer (CL-500A, KONICA MINOLTA, Inc., Japan), and spectral power distributions under Light conditions are shown in Fig. S1. Melanopic equivalent daylight illuminance (EDI) of the light conditions was 155.1 melanopic lx. The information on light stimulus is reported as the ENLIGHT Checklist and Guidelines^27^

Saliva samples were collected hourly from 17:00 using Salivette cotton swabs (Sarstedt AG & Co. KG, Germany). In the Pre-camp session, sampling continued until each child’s habitual bedtime, and the final sample was obtained between 20:00 and 23:00 (mean final sampling time, approximately 22:00 ± 0.8 hours), while the sampling continued until 20:00 for all participants in the Camp session (Fig. 1). During sampling, participants engaged in quiet activities such as reading, studying, or conversation, except during saliva sampling. Participants were instructed to finish dinner by 17:00 on the day of melatonin measurements and to sit in a chair until the saliva sample collection procedure was complete. Participants did not drink any water for 15 min prior to each salivary sample collection. When brief outdoor movement from the shed was unavoidable, such as going to the restroom, participants wore sunglasses to limit uncontrolled light exposure.

### Melatonin assay, DLMO, and suppression

Salivary melatonin concentrations were quantified using a radioimmunoassay kit (RK-DSM2, Bühlmann Laboratories AG, Switzerland). Dim light melatonin onset (DLMO), which is a marker of circadian phase^28,29^, was determined from the melatonin profile under the dim-light condition using a fixed threshold of 4.0 pg/mL, a threshold commonly used in pediatric and adolescent studies^30,31^. The time of threshold crossing was estimated by linear interpolation between the samples immediately below and above 4.0 pg/mL. Participants whose dim-light melatonin concentration did not exceed 4.0 pg/mL before the final sample were excluded from analyses requiring DLMO.

For each condition, i.e., Light and Dim conditions, the area under the curve (AUC) of the melatonin concentration time course curve was calculated from the DLMO to the final sampling time using linear trapezoidal integration. For the light condition, integration began at the clock time corresponding to the DLMO obtained under dim-light conditions. Melatonin suppression was calculated as: suppression (%) = [(*AUCdim* – *AUClight*) / *AUCdim*] × 100. Negative values indicated a larger AUC under light conditions than under dim-light conditions.

### Habitual sleep timing

Habitual sleep variables were derived from the 14–day baseline sleep diaries. As mentioned above, sleep-related data obtained from wrist activity monitors were excluded from further analysis because they were frequently missing. Sleep onset was calculated from reported bedtime and sleep-onset latency, wake time was the reported final awakening time, sleep duration was the interval from sleep onset to wake time, and sleep midpoint was the midpoint of that interval. Participant-level values were averaged across available diary days.

### Personal light-exposure assessment

Personal light exposure was recorded as wrist-level photopic illuminance at 1-min intervals using the light sensor incorporated in the MotionWatch 8. As mentioned above, light exposure data obtained from wrist activity monitors before the camping program were not available because they were frequently missing. Therefore, the light-exposure data were analysed and used only in the Camp session.

In the current study, two complementary exposure metrics were calculated: geometric mean illuminance, representing typical sustained exposure, and cumulative illuminance (illuminance dose), representing the combined intensity and duration of exposure^32,33^. To account for the strongly skewed distribution and zero values in illuminance data, geometric mean illuminance was calculated by averaging the epoch-level values after transformation using log10(lx + 0.1) and then back-transforming the results^34^. Illuminance dose was accumulated from the untransformed illuminance values and expressed in lx-hours.

For the main exploratory analyses, both metrics were calculated from 5:00 to 17:00 on the corresponding Camp Light-condition day, thereby excluding the experimental light exposure beginning at 17:00. As an additional exploratory assessment of prior light history, daily values were calculated from 5:00 to 20:00 during the seven days preceding the Light-condition day in the Camp session and then averaged within each participant. An analysis window was considered valid when at least 80% of the expected measurements were available. The 7-day exposure metrics were calculated when at least six of the seven days were valid.

### Statistical analysis

Statistical analysis was performed using R^35^ version 4.6.1. Associations of Pre-camp melatonin suppression with Pre-camp DLMO and habitual sleep variables (bedtime, sleep onset time, wake time, sleep duration and mid-sleep time) were examined using Spearman’s rank correlation coefficients among participants with available paired data.

Associations between personal light exposure and melatonin suppression were also examined using Spearman’s rank correlation coefficients in only the Camp session. The main exploratory analyses used light exposure measured from 5:00 to 17:00 on the corresponding Light-condition day. Additional exploratory analyses used the participant-level exposure values averaged over the seven days preceding the Light-condition day.

Sensitivity analyses were conducted to assess the influence of outlier values. For the associations of Pre-camp suppression with DLMO and habitual sleep variables, the analyses were repeated after excluding outliers in Pre-camp melatonin suppression. For the light exposure analyses, the correlations were repeated after excluding participants with an outlying value in the Camp melatonin suppression. For light exposure metrics, illuminance dose and geometric mean illuminance, outliers were identified separately for each metric. This exclusion based on IQR for melatonin suppression was also applied when comparing melatonin suppression between Pre-camp and Camp sessions. Outliers were defined using the interquartile range (IQR) criterion as values below *Q*_1_ – 1.5 × IQR or above *Q*_3_ + 1.5 × IQR, where *Q*_1_ and *Q*_3_ represent the first and third quartiles, respectively, and IQR = *Q*_3_ – *Q*_1._

As a post hoc complementary exploratory analysis, participants were divided into lower- and higher-suppression groups using mean value of Pre-camp melatonin suppression as the cut-off. Participants with suppression below mean value were classified as the lower-suppression group, whereas those with suppression of mean value or greater were classified as the higher-suppression group. DLMO and habitual sleep variables were compared between the groups using Welch’s two-sample *t*-test.

Melatonin suppression in the Pre-camp and Camp sessions was compared among participants with valid values in both periods using a paired-samples *t*-test. We used the Pitman–Morgan test to compare variability in melatonin suppression between the Pre-camp and Camp sessions. All tests were two-sided, with a statistical significance level of *p* < 0.05. Given the exploratory and hypothesis-generating nature of the study, no adjustment was made for multiple comparisons.

## Results

### Individual variability in melatonin suppression

Pre-camp and Camp melatonin suppression were available for 20 and 19 children, respectively. One participant in the Pre-camp session and two participants in the Camp session were excluded because the increase in melatonin under the Dim condition did not exceed the 4.0 pg/mL DLMO threshold used to calculate melatonin suppression. The individual melatonin profiles in the Pre-camp and Camp session are shown in Fig. S2 and S3, respectively. As previously reported^26^, mean DLMO occurred at 20:50 ± 0:45 at the Pre-camp assessment and at 18:43 ± 0:20 at the Camp assessment. The mean within-participant advance was 2:06 ± 0:45 (n = 15, paired *t*-test: *p* < 0.001. Suppression values were widely distributed among participants, ranging from negative values to marked suppression in both sessions (Fig. 2). For the Pre-camp melatonin suppression, the mean ± SD of Pre-camp melatonin suppression was 39.0 ± 52.6 and twelve children had suppression values of mean value or greater, whereas eight had values below mean value. The distributions included one extreme negative value in the Pre-camp session and two outliers in the Camp session identified using the IQR. After excluding these values, substantial interindividual variability remained, especially in the Pre-camp session (Fig. 2).

**Figure 2.**
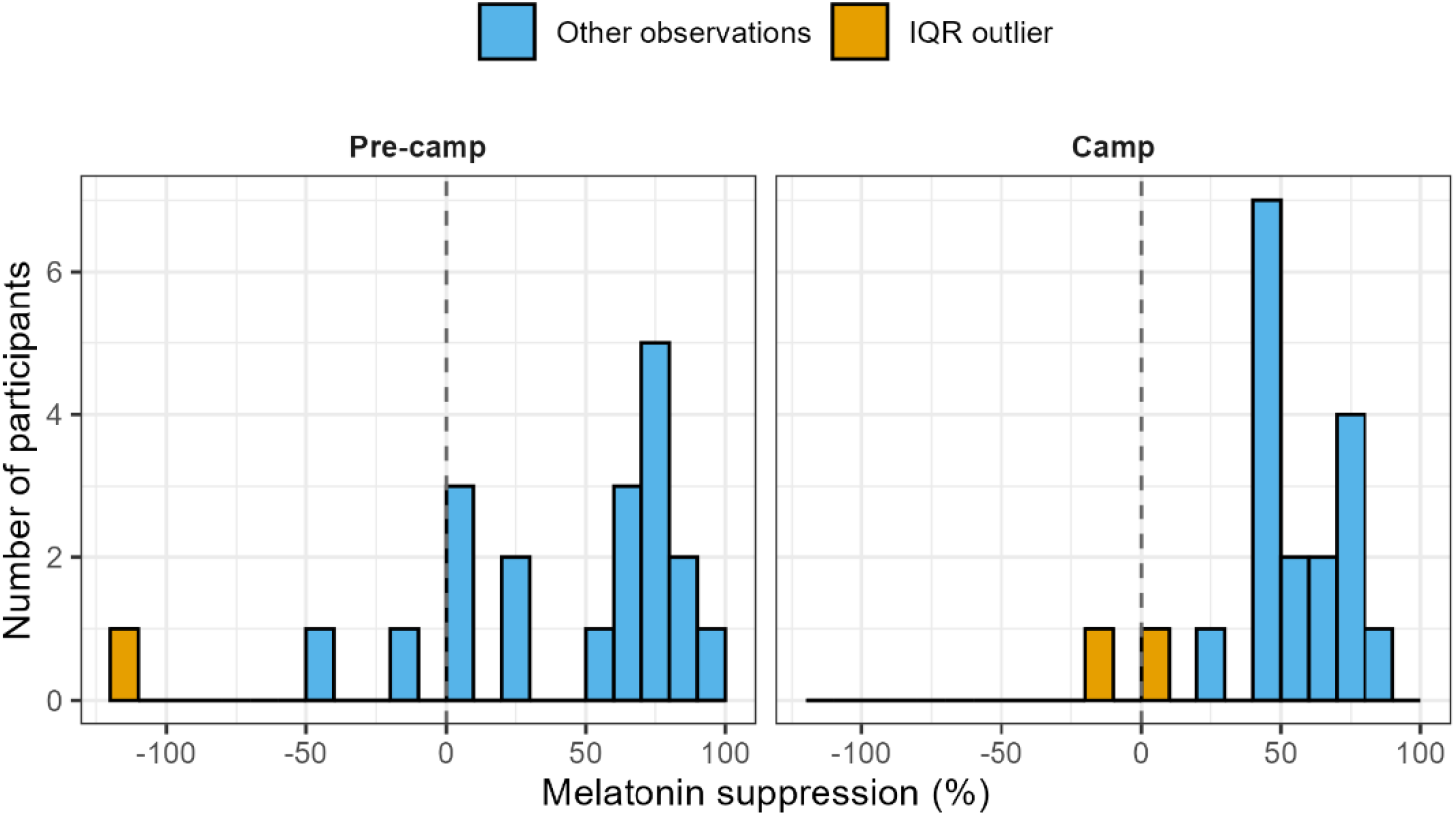
Distribution of melatonin suppression in Pre-camp and Camp sessions. Orange bars indicate observations identified as outliers in melatonin suppression using the IQR criterion.

### Comparison in melatonin suppression between Pre- and Camp sessions

Paired Pre-camp and Camp session melatonin suppression values were available for 18 children. Mean melatonin suppression was 34.8 ± 53.9% in the Pre-camp session and 51.1 ± 25.1% in the Camp session (Fig. 3). The mean within-participant change was 16.3 percentage points, but the confidence interval was wide and included zero (95% CI −10.2 to 42.8 percentage points). The difference between the Pre-camp and Camp assessments was not statistically significant (paired *t*(17) = 1.30, *p* = 0.211). After excluding participant pairs containing an IQR outlier in either session, the difference remained non-significant (n = 15, paired *t*(14) = 1.42, *p* = 0.177). Although melatonin suppression did not differ significantly between sessions, its distribution in the Camp session was significantly narrower than in the Pre-camp session (Pitman–Morgan test: *t*(16) = 3.47, *p* = 0.003).

**Figure 3.**
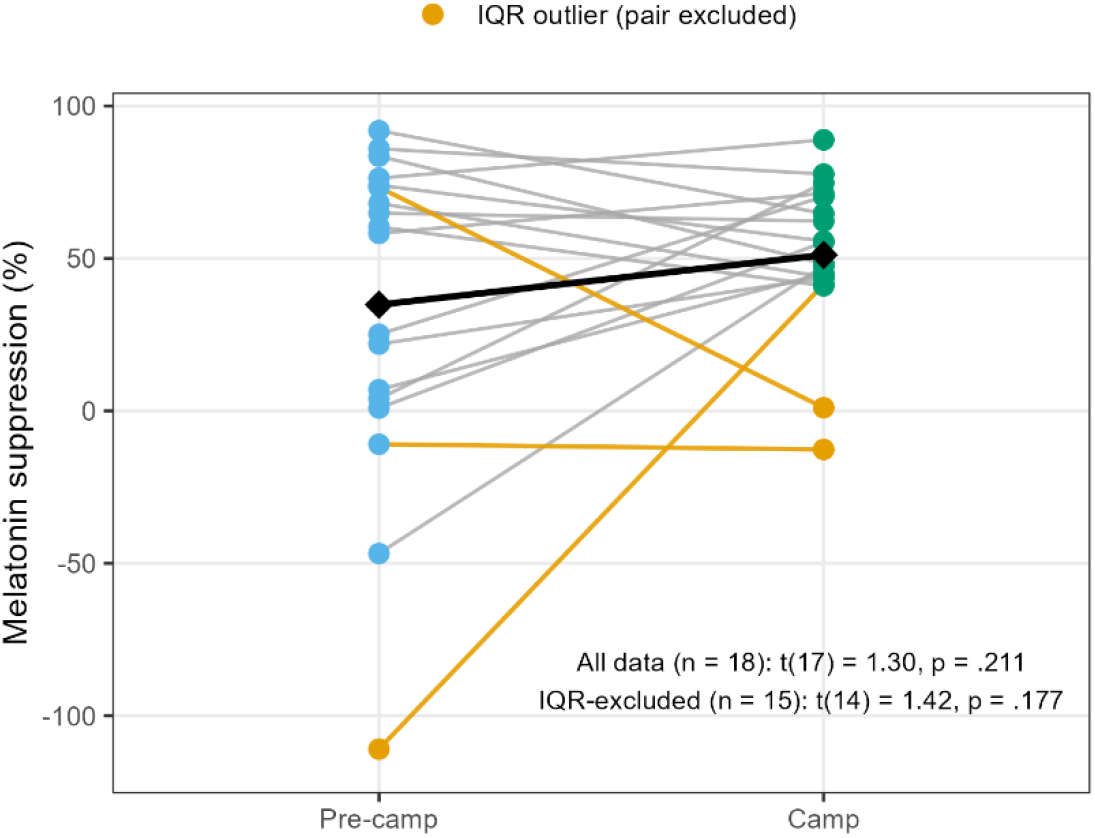
Comparison of melatonin suppression before and during camp. Each line connects observations from the same participant. Blue and green circles indicate non-outlying observations before and during camp, respectively. Orange circles indicate observations identified as outliers using the IQR criterion, applied separately within each session. Orange lines identify participant pairs containing an outlier in either session, and these pairs were excluded from the IQR-excluded sensitivity analysis. Black diamonds and the connecting black line represent mean values calculated from all paired observations.

### Associations of Pre-camp melatonin suppression with DLMO and habitual sleep variables

Among the twenty participants with analyzable melatonin suppression data, three participants were excluded from the examination of associations with habitual sleep variables because their sleep diary data were frequently missing. Greater Pre-camp melatonin suppression was associated with later DLMO (Spearman’s *rho* = 0.49, *p* = 0.028). For reference, the correlation between DLMO and melatonin suppression in the Camp-session (*rho* = 0.31, *p* = 0.20, *n* = 19) and the correlation between changes in DLMO and changes in melatonin suppression from Pre-camp to Camp were not significant (*rho* = 0.29, *p* = 0.25, *n* = 18). For the habitual sleep measures, melatonin suppression was not significantly associated with bedtime (*rho* = 0.37, *p* = 0.15), sleep onset time (*rho* = 0.40, *p* = 0.11), wake time (*rho* = 0.31, *p* = 0.22), midsleep time (*rho* = 0.34, *p* = 0.18) and sleep duration (*rho* = −0.19, *p* = 0.47; Fig. 4 and Table 1). Melatonin suppression was not significantly associated with either the duration of experimental light exposure from 17:00 to the final sampling time, i.e., habitual bedtime (*rho* = 0.31, *p* = 0.19) or the AUC integration interval from DLMO to the final sampling time (*rho* = −0.32, *p* = 0.16). To assess whether the correlation between DLMO and melatonin suppression depended on the length of the AUC integration window, we recalculated melatonin suppression using a fixed 30-min window beginning at DLMO. Among the 17 participants with sufficient data in both conditions, the correlation was similar for the original time window, i.e., DLMO to final sampling time (*rho* = 0.44, *p* = 0.08), and the fixed 30-min windows (*rho* = 0.44, *p* = 0.08).

**Figure 4.**
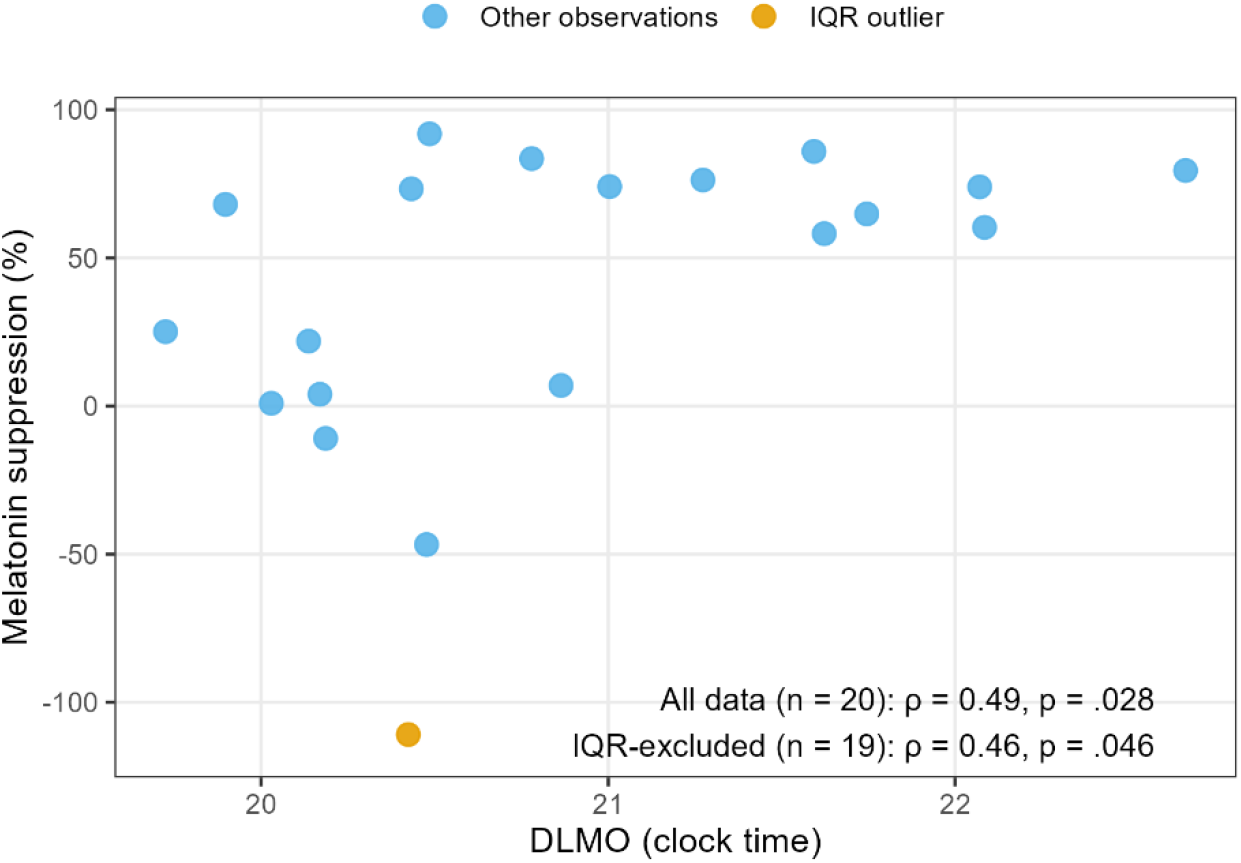
Relationship between Pre-camp melatonin suppression and DLMO. Orange circles indicate observations identified as outliers in melatonin suppression using the IQR criterion.

**Table 1.**
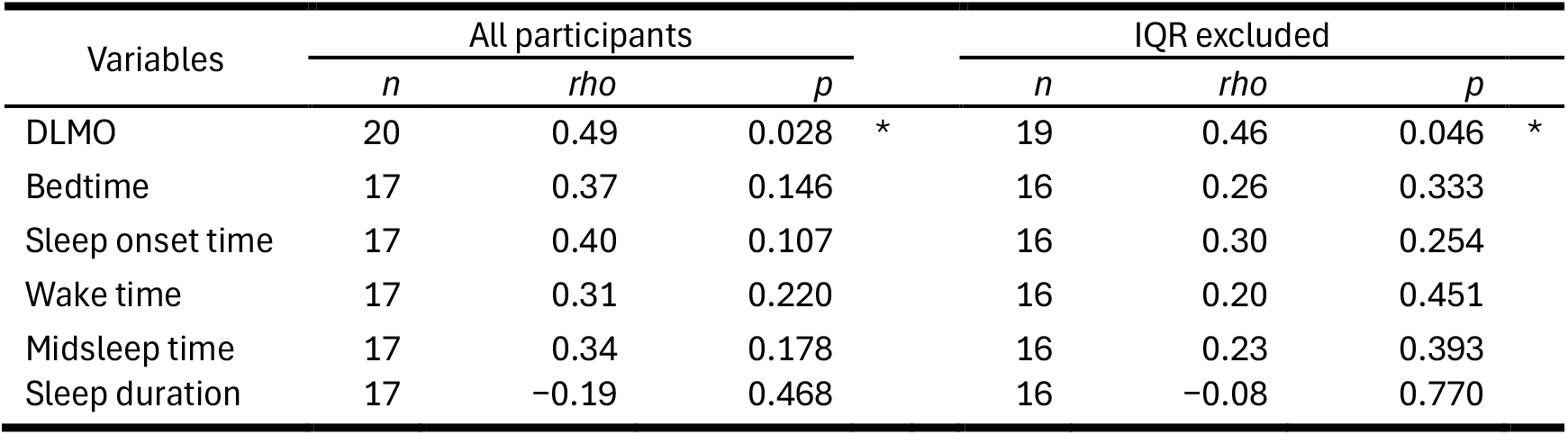
Relationships between Pre-camp melatonin suppression and circadian phase: DLMO or habitual sleep variables. IQR excluded represents the sample size and statistical values after excluding observations of melatonin suppression identified as outliers. *: *p* < 0.05.

| Variables | All participants |  |  |  | IQR excluded |  |  |  |
|---|---|---|---|---|---|---|---|---|
|  | *n* | *rho* | *p* |  | *n* | *rho* | *p* |  |
| DLMO | 20 | 0.49 | 0.028 | * | 19 | 0.46 | 0.046 | * |
| Bedtime | 17 | 0.37 | 0.146 |  | 16 | 0.26 | 0.333 |  |
| Sleep onset time | 17 | 0.40 | 0.107 |  | 16 | 0.30 | 0.254 |  |
| Wake time | 17 | 0.31 | 0.220 |  | 16 | 0.20 | 0.451 |  |
| Midsleep time | 17 | 0.34 | 0.178 |  | 16 | 0.23 | 0.393 |  |
| Sleep duration | 17 | −0.19 | 0.468 |  | 16 | −0.08 | 0.770 |  |

To assess the influence of extreme suppression values, the analyses were repeated after excluding the value identified by the IQR criterion. The positive association between melatonin suppression and DLMO remained (*rho* = 0.46, *p* = 0.046), whereas none of the associations with the habitual sleep variables was statistically significant (all *p* > 0.1).

### Exploratory comparisons between higher- and lower-suppression groups

As a complementary exploratory analysis, participants were divided into higher-(≥ mean value) and lower-suppression (< mean value) groups. DLMO was later in the higher-suppression group than in the lower-suppression group (21:18 ± 0:49 h vs 20:15 ± 0:20 h; mean difference, 1.05 h; 95% CI 0.50 to 1.61 h; Welch’s *t*(15.84) = 4.01, *p* = 0.001: Fig. S4).

The higher-suppression group also had a later habitual bedtime (22:29 ± 0:45 h vs. 21:26 ± 0:47 h; mean difference, 1.05 h; 95% CI 0.24 to 1.87 h; *p* = 0.015), later sleep onset (22:40 ± 0:48 h vs. 21:32 ± 0:45 h; mean difference, 1.13 h; 95% CI 0.32 to 1.95 h; *p* = 0.010), and later midsleep time (02:47 ± 0:44 h vs. 01:50 ± 0:41 h; mean difference, 0.94 h; 95% CI 0.20 to 1.68 h; *p* = 0.017). Differences in wake time (*p* = 0.068) and sleep duration (*p* = 0.216) were not statistically significant (Fig. S4).

### Associations of Camp melatonin suppression with personal light exposure

Valid paired melatonin suppression and light exposure data for the experimental-day metrics were available for 15 participants in the Camp session. For light exposure measured from 5:00 to 17:00 on the corresponding Light-condition day, greater same-day geometric mean illuminance was associated with lower melatonin suppression (*rho* = −0.58, *p* = 0.024). The association with illuminance dose was also negative but did not reach statistical significance (*rho* = −0.38, *p* = 0.160; Fig. 5 and Table 2).

**Figure 5.**
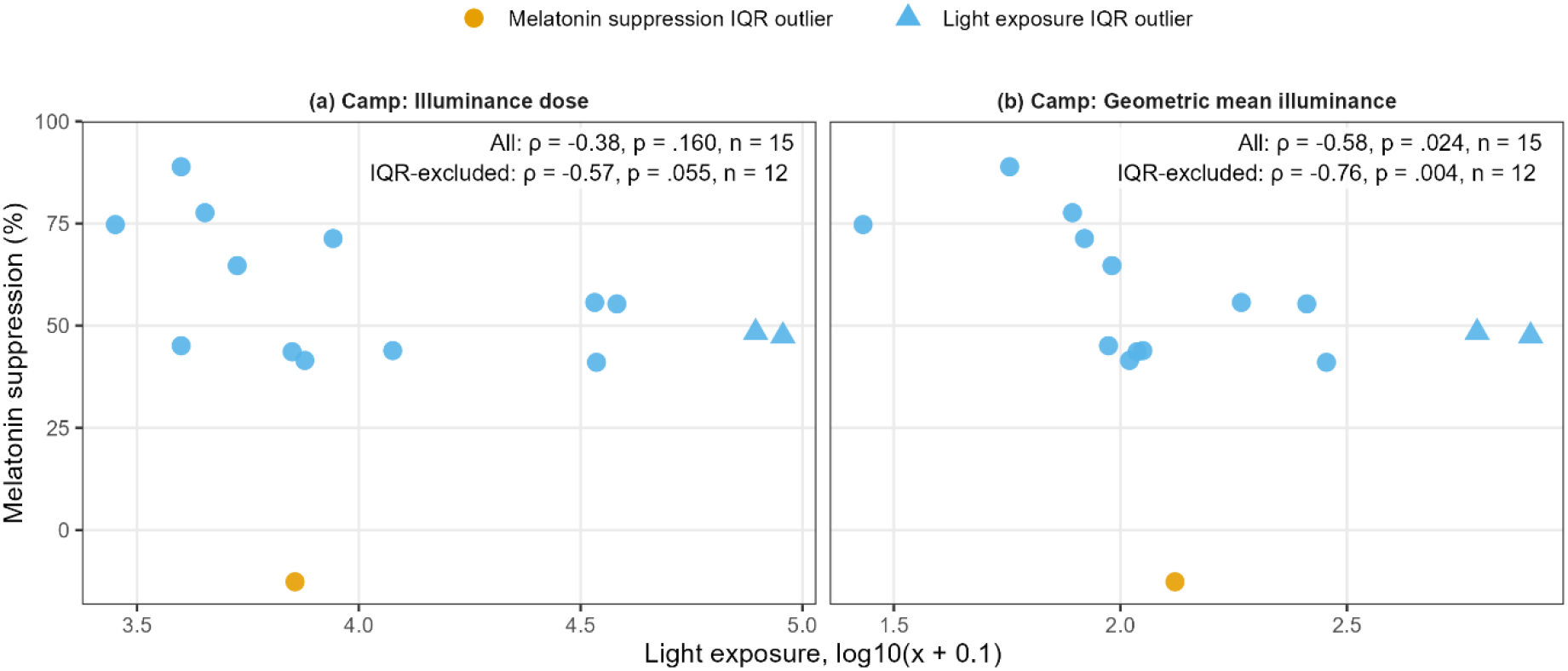
Relationships between melatonin suppression and light exposure metrics in the Camp session. (a) Illuminance dose, and (b) geometric mean illuminance on the day conducted the Light condition. Orange circles and blue triangles indicate observations identified as outliers using the IQR criterion in melatonin suppression and light exposure metrics, respectively.

**Table 2.** Relationships between melatonin suppression and light exposure metrics: illuminance dose and the geometric mean of the experimental day (Light condition), or the 7-day mean preceding the experimental day in the Camp session. *: *p* < 0.05, **: < 0.01.

| Exposure window | Exposure metrics | All participants |  |  |  | IQR excluded |  |  |  |
|---|---|---|---|---|---|---|---|---|---|
|  |  | *n* | *rho* | *p* |  | *n* | *rho* | *p* |  |
| Experimental day | Dose | 15 | −0.38 | 0.160 |  | 12 | −0.57 | 0.055 |  |
|  | Geometric mean | 15 | −0.58 | 0.024 | * | 12 | −0.76 | 0.004 | ** |
| Previous 7-day | Dose | 15 | −0.37 | 0.178 |  | 13 | −0.32 | 0.289 |  |
|  | Geometric mean | 15 | −0.24 | 0.383 |  | 13 | −0.30 | 0.325 |  |

In the additional analyses of light exposure averaged over the preceding seven days, no statistically significant associations were observed. Neither association was statistically significant for either illuminance dose (*rho* = −0.37, *p* = 0.177; *n* = 15) or geometric mean illuminance (*rho* = −0.24, *p* = 0.383; *n* = 15).

Sensitivity analyses excluding IQR-defined outliers in melatonin suppression or the corresponding light-exposure variable produced broadly similar results. The negative association between same-day geometric mean illuminance and Camp melatonin suppression remained (*rho* = −0.76, *p* = 0.004; *n* = 12). The corresponding association with illuminance dose remained negative but was not statistically significant (*rho* = −0.57, *p* = 0.055; *n* = 12). None of the associations based on the preceding seven days was statistically significant after combined IQR exclusion (all *p* ≥ 0.289).

## Discussion

This exploratory study showed interindividual variability in light-induced melatonin suppression among school-aged children. The planned comparison between the Pre-camp and Camp sessions did not provide clear evidence of a mean change in melatonin suppression. For the interindividual variability in melatonin suppression, greater Pre-camp melatonin suppression was associated with later DLMO, whereas associations with habitual sleep variables were not statistically significant. In the Camp session, greater geometric mean illuminance on the day of the Light-conditions experiment was associated with lower melatonin suppression that evening, although no associations were found for light exposure averaged over the preceding seven days. These findings did not provide clear evidence of a change in mean melatonin suppression across the camping period, but suggest that circadian timing and recent daytime light exposure may be associated with children’s light-induced melatonin suppression.

Melatonin suppression before camping varied widely among children, ranging from negative values to greater suppression. The previous study examining the school-aged children’s melatonin suppression reported marked melatonin suppression in children exposed to 580 lx light^15^. The lower illuminance used in the present study (200 lx) may have allowed greater variability in suppression to be observed, whereas responses at higher illuminance may have been closer to a ceiling. However, differences in other experimental conditions also limit direct comparison between the studies. On the other hand, our finding is consistent with the large interindividual variability reported in adults. Phillips et al. observed a more than 50-fold difference among individuals in the illuminance required to produce 50% melatonin suppression^18^. However, unlike the dose-response approach used by Phillips et al., suppression was assessed at only one light level and once per session in the present study. The observed variability therefore cannot be attributed specifically to stable individual differences in light sensitivity and may also reflect differences in circadian timing, melatonin secretion profiles, or measurement variability. Repeated assessments across multiple light levels are needed to determine whether school-aged children show reproducible individual differences in melatonin suppression.

We explored factors potentially associated with melatonin suppression variability using the data available in each session. The Pre-camp session was conducted before the camping schedule modified the children’s habitual circadian phase and sleep timing. In this session, greater melatonin suppression was associated with later DLMO, indicating that children with a later circadian phase showed a greater suppression response by evening light exposure. A similar relationship has been reported by Moderie et al., who found that melatonin suppression was positively associated with circadian phase within adults with a delayed sleep schedule^20^. One possible explanation is that children who respond more strongly to evening light experience greater suppression and phase-delaying effects from habitual evening illumination, thereby contributing over time to a later circadian phase. Meanwhile, the association between later DLMO and greater melatonin suppression requires cautious interpretation. In this session, we set the start and end times of light exposure at 17:00 and each participant’s habitual bedtime. Consequently, the participants with later sleep habits might have been exposed for a longer duration than those with earlier habits. Since light exposure duration is the factor affecting melatonin suppression^36^, there is a possibility that the relationship between light exposure duration and later sleep habits leads to an overestimation of the association of melatonin suppression with DLMO. In addition, DLMO was used to define the beginning of the AUC interval from which suppression was calculated. Although neither the duration of experimental light exposure nor the DLMO-to-final-sample integration interval was significantly associated with melatonin suppression, mathematical dependence on DLMO and differences in the rising melatonin profile cannot be excluded. Future studies specifically designed to test this hypothesis are therefore required.

In the complementary group analysis, children in the higher-suppression group also had later habitual bedtime, sleep onset and midsleep time than those in the lower-suppression group. This pattern is broadly consistent with the possibility proposed by a previous study^19^, which observed earlier habitual bedtimes in adults showing little or no melatonin suppression. However, the continuous correlations with habitual sleep variables were not statistically significant in the present study and were attenuated after exclusion of the IQR-defined suppression outlier. Moreover, the mean value threshold (39.0%) used for the group analysis was exploratory. The group findings should therefore be considered hypothesis-generating rather than confirmatory evidence of an association between habitual sleep timing and melatonin suppression.

During the camping, we explored whether naturally occurring daytime light exposure preceding the controlled evening-light experiment was associated with melatonin suppression. Greater same-day geometric mean illuminance was associated with lower melatonin suppression, while illuminance dose showed a negative association in the same direction but did not reach statistical significance. These findings suggest that children exposed to higher daytime geometric mean illuminance during the day may show a lower melatonin suppression response to light that evening. This direction is consistent with controlled studies showing that prior exposure to relatively bright light attenuates subsequent melatonin suppression, whereas a dimmer light history increases the response^22,23^. Seasonal observations have similarly demonstrated greater melatonin suppression following lower daytime light exposure in winter^24^. In addition, our result is consistent with the field study showing the negative correlation between melatonin suppression and averaged 24-h light exposure observed in indoor and outdoor workers^37^. The present findings extend this possibility to naturalistic daytime light exposure during camping in children. However, these findings should be interpreted cautiously because of the small number of participants with valid light-exposure data and the exploratory and hypothesis-generating analysis. Therefore, the association should be examined in future studies with prospectively defined exposure windows and more complete personal light measurements.

The planned comparison did not detect a statistically significant difference in mean melatonin suppression between the Pre-camp and Camp sessions. Similarly, light exposure averaged over the preceding seven days was not associated with suppression during camping. Neither the Pre-camp versus Camp comparison nor the exploratory correlations with seven-day light exposure provided clear evidence of an association with melatonin suppression. Hébert et al. demonstrated that melatonin suppression in adults was lower after a week of increased daytime bright-light exposure (bright week) than after a week of restricted light exposure (dim week)^38^, and our findings may seem to contradict this result. However, whereas the previous study increased or restricted daytime light exposure, the present study compared habitual living conditions before camping with the natural light environment during camping. The contrast in prior light exposure may therefore have been smaller than in the previous study, potentially resulting in a smaller effect on melatonin suppression. Melatonin suppression varied less between children in the Camp session than in the Pre-camp session. If circadian timing and prior light exposure contribute to individual differences in melatonin suppression, a more standardised sleep schedule, less variation in circadian phase, similar daily activities and light exposure during camp may have contributed to the narrower distribution. Indeed, DLMO variation in the Camp session (18:43 ± 0:20) was significantly lower than in the Pre-camp session (20:50 ± 0:45), not only advanced it^26^. Conversely, the difference in melatonin suppression variation may have resulted from differences in the experimental protocols or from the influence of extreme Pre-camp values. Therefore, it should be regarded only as a hypothesis for future investigation.

This study has several limitations. First, the sample size was small and consisted predominantly of boys, and missing sleep habits and/or light-exposure data further reduced the sample available for some analyses. We could not analyse the sleep habits and light exposure data from wrist-worn loggers before the camping program because they were frequently missing: participants could not wear them at school. Therefore, sleep habits data before the camping were assessed by self-reported sleep diary, and the increased light exposure in the Camp session remained hypothetical. Multiple exploratory analyses were conducted without adjustment for multiple comparisons. Unlike previous studies characterising individual sensitivity using dose-response curves^18^, suppression was assessed at only one light level and once per session. The protocols also differed between the Pre-camp and Camp sessions in exposure and sampling duration and in the order of the Light and Dim conditions. In addition, the Dim condition of < 30 lx may itself have affected melatonin secretion in these light-sensitive children, but we used incandescent lamps and melanopic EDI is predicted to be around 15 according to the CIE S 026 Toolbox^39^. Finally, daytime light exposure was measured as wrist-level photopic illuminance, which does not directly represent light reaching the eye and may be affected by sensor orientation, clothing and local shading.

In conclusion, school-aged children showed wide variability in their melatonin suppression response to evening light in the specific setting of a camping program. Later DLMO and lower same-day daytime light exposure were identified as potential correlates of greater melatonin suppression in children, while the exploratory group comparison generated an additional hypothesis concerning habitual sleep timing. Despite its limitations, this study provides hypotheses that circadian phase and recent light history may contribute to children’s responses to evening light outside highly controlled laboratory settings. Future studies with larger samples, repeated assessments and prospectively defined personal light-exposure measurements are needed to test these hypotheses.

## Supporting information

ENLIGHT Checklist

## Acknowledgements

The authors would like to thank Ryuichiro Sakino, producer of the camping program, and all camping staff in Mobilityland Corporation in Japan for their understanding and cooperation in our study. During the preparation of this manuscript, the authors used ChatGPT (OpenAI) to support English-language editing and improve clarity, and to develop and debug R code. The authors critically reviewed, verified, and revised all AI-assisted outputs. The authors take full responsibility for the content of the manuscript, the analysis plan, and the accompanying code.

## Author contributions

TE, KT, KN, MO, SL, SK, AS, SN and SH contributed to the design of the experiment. TE, KT, KN, MO, SL, YI, YN, SK and SH performed the experiment. TE, KN, KT and MO analysed the data. TE and KT wrote the manuscript. KN, MO, SL, YI, YN, SK, AS, SN and SH participated in the discussion and preparation of the manuscript. All authors read and approved the final manuscript.

## Statements and declarations

### Ethical considerations

The Ethical Committee of Kyushu University approved this study before it was conducted. The experiments were conducted in accordance with the Declaration of Helsinki.

### Consent to participate

Written informed consent was obtained from the parents of all participating children after confirming the children’s willingness to participate.

### Data availability

The datasets analysed in this study are not publicly available to protect participant privacy but are available from the corresponding author upon reasonable request.

