## Supplementary material for "Light-induced melatonin suppression in school-aged children before and during camping: associations with circadian phase and daytime light exposure": ENLIGHT Checklist

Note: This document is best viewed and completed with Adobe Acrobat Reader.

ENLIGHT

Release 1.0.2, 16 October 2023

### ENLIGHT Checklist

Below is the **ENLIGHT Checklist** for reporting ocular light exposures in human laboratory-based studies. We will strongly encourage that this checklist be used in conjunction with the **ENLIGHT Explanation & Elaboration (E&E) document**. This checklist is intended both to help authors, reviewers, and editors in evaluating the completeness of reporting in submitted studies, and for documentation of studies after publication. In the location column, please indicate the page, figure, or table number where the item or description can be found. If an item is not available, please select "**Not available**". If you consider an item not to be applicable in your specific study design after consulting the guidelines, please select "**Not applicable**". Items which do not have the option to select "Not applicable" were rated by experts as applicable for all studies, regardless of context. If you are unable to provide the information, please select "Not available".

The **ENLIGHT Checklist** (this document) and the **ENLIGHT E&E document** are released under the CC-BY-NC-ND License. For more information, please visit http://enlight-statement.org/.

#### General Information

**Author names:** See appendix: authors

**Title of manuscript:** See appendix: title

**Date:** 2026-11-12

#### A. Study Characteristics

##### A.1. Protocol-level characteristics

| | Location (page, figure, table number) | Not available | Not applicable |
|---|---|---|---|
| Description of experimental setting | Methods, pp. 6-8. | ☐ | |
| Timeline of experiment (including timing and duration of light) | Methods, pp. 6-8, Figure 1. | ☐ | |
| Pre-laboratory sleep-wake/rest-activity behaviour | Methods, pp. 6 and 8 | ☐ | ☐ |
| Pre-laboratory light exposure | Methods, p. 9 | ☐ | ☐ |
| Immediate prior light exposure (in laboratory) | | ☑ | ☐ |

##### A.2. Measurement-level characteristics

| | | | |
|---|---|---|---|
| Measurement plane (e.g., horizontal or vertical) | Methods, p. 7 | ☐ | |
| Measurement viewpoint and location | Methods, p. 7 | ☐ | |
| Type, make and manufacturer of the measurement instrument | Methods, p. 7 | ☐ | |
| Calibration status of the instrument | | ☑ | ☐ |

##### A.3. Participant-level characteristics

| | | | |
|---|---|---|---|
| Ocular health and functioning | | ☑ | |
| Pupil size and/or dilation | | ☑ | ☐ |
| Relative time (e.g. to circadian phase or sleep) | Methods, pp. 6-7 | ☐ | ☐ |

### B. Light characteristics

#### B.1. Light source type(s). Please select all that are relevant.

| Room illumination (overhead or other) | Emissive surfaces including displays (incl. light therapy devices) | Wearable light emitting glasses | Ganzfeld exposure | Other: |
|---|---|---|---|---|
| ☑ | ☐ | ☐ | ☐ | ☐ |

| Polychromatic light | Monochromatic or narrowband light |
|---|---|
| ☑ | ☐ |

| | Location (page, figure, table number) | Not available | Not applicable |
|---|---|---|---|
| Type, make and manufacturer of the light source | Methods, p. 7 | ☐ | |
| Use of wearable filtering apparatus (e.g., blue-blocking glasses) | Methods, p. 8 | ☐ | ☐ |

#### B.2. Light level characteristics

| | | | |
|---|---|---|---|
| Illuminance (lux) and/or luminance (cd/m$^2$) | Methods, p. 7 | ☐ | |
| Spectral irradiance and/or radiance distribution | Supplementary Figure S1 | ☐ | ☐ |
| α−opic irradiance and/or radiance (including melanopic) | | ☑ | ☐ |
| α−opic equivalent daylight illuminance and/or luminance (EDI/EDL, including melanopic) | Methods, p. 7 | ☐ | ☐ |

**NOTE:** Luminance and radiance metrics (as opposed to illuminance and irradiance) are mainly relevant for emissive surfaces.

#### B.3. Colour characteristics

| | | | |
|---|---|---|---|
| Peak wavelength and bandwidth | See appendix: B3.1 | ☐ | ☐ |
| Colour appearance quantities (any) | Methods, p. 7 | ☐ | ☐ |
| Colour rendering metrics (any) | | ☑ | ☐ |

**NOTE:** Peak wavelength and bandwidth are most relevant for monochromatic or narrowband light sources.

#### B.4. Temporal and spatial characteristics

| | | | |
|---|---|---|---|
| Location of stimulus and viewing distance | Methods, p. 8 | ☐ | |
| Temporal pattern (including flash frequency and waveform) | Methods, p. 7; Figure 1 | ☐ | ☐ |
| Relative or absolute size of the stimulus | | ☑ | ☐ |

Reset form to default values    Print as PDF

### Completed response record

ENLIGHT Checklist v1.0.2 | 25 of 25 items addressed

This appendix records the information entered in the ENLIGHT wizard. The preceding two pages retain the official checklist. Item-level identifiers below are wizard identifiers; the official checklist numbers sections only.

#### Manuscript

Title: Light-induced melatonin suppression in school-aged children before and during camping: associations with circadian phase and daytime light exposure

Authors: Taisuke Eto, Kota Takeoka, Kana Nishimura, Michihiro Ohashi, Sang-il Lee, Yuki Ikeda, Yuki Nishimura, Shingo Kitamura, Akiko Shikano, Shingo Noi, Shigekazu Higuchi

Date: 2026-11-12

#### B1.1 Light source type(s)

Source types: Room illumination (overhead or other)

Spectral types: Polychromatic light

#### A.1. Protocol-level characteristics

##### A1.1 Description of experimental setting

Status: Reported

Location: Methods, pp. 6-8.

##### A1.2 Timeline of experiment (including timing and duration of light)

Status: Reported

Location: Methods, pp. 6-8, Figure 1.

##### A1.3 Pre-laboratory sleep-wake/rest-activity behaviour

Status: Reported

Location: Methods, pp. 6 and 8

##### A1.4 Pre-laboratory light exposure

Status: Reported

Location: Methods, p. 9

Notes: Only Camp session data

##### A1.5 Immediate prior light exposure (in laboratory)

Status: Not available

#### A.2. Measurement-level characteristics

##### A2.1 Measurement plane (e.g., horizontal or vertical)

Status: Reported

Location: Methods, p. 7

##### A2.2 Measurement viewpoint and location

Status: Reported

Location: Methods, p. 7

##### A2.3 Type, make and manufacturer of the measurement instrument

Status: Reported

Location: Methods, p. 7

##### A2.4 Calibration status of the instrument

Status: Not available

#### A.3. Participant-level characteristics

##### A3.1 Ocular health and functioning

Status: Not available

##### A3.2 Pupil size and/or dilation

Status: Not available

##### A3.3 Relative time (e.g. to circadian phase or sleep)

Status: Reported

Location: Methods, pp. 6-7

#### B.1. Light source type(s)

##### B1.2 Type, make and manufacturer of the light source

Status: Reported

Location: Methods, p. 7

##### B1.3 Use of wearable filtering apparatus (e.g., blue-blocking glasses)

Status: Reported

Location: Methods, p. 8

#### B.2. Light level characteristics

##### B2.1 Illuminance (lux) and/or luminance (cd/m²)

Status: Reported

Location: Methods, p. 7

##### B2.2 Spectral irradiance and/or radiance distribution

Status: Reported

Location: Supplementary Figure S1

##### B2.3 α−opic irradiance and/or radiance (including melanopic)

Status: Not available

##### B2.4 α−opic equivalent daylight illuminance and/or luminance (EDI/EDL, including melanopic)

Status: Reported

Location: Methods, p. 7

Notes: Melanopic EDI only; Spectral distribution is reported

#### B.3. Colour characteristics

##### B3.1 Peak wavelength and bandwidth

Status: Reported

Location: Supplementary Figure S1 (as a spectral distribution)

##### B3.2 Colour appearance quantities (any)

Status: Reported

Location: Methods, p. 7

Notes: CCT

##### B3.3 Colour rendering metrics (any)

Status: Not available

#### B.4. Temporal and spatial characteristics

##### B4.1 Location of stimulus and viewing distance

Status: Reported

Location: Methods, p. 8

##### B4.2 Temporal pattern (including flash frequency and waveform)

Status: Reported

Location: Methods, p. 7; Figure 1

##### B4.3 Relative or absolute size of the stimulus

Status: Not available

#### Guideline source

Spitschan M, Kervezee L, Lok R, McGlashan E, Najjar RP; ENLIGHT Consortium. ENLIGHT: A consensus checklist for reporting laboratory-based studies on the non-visual effects of light in humans. EBioMedicine. 2023;98:104889.

https://doi.org/10.1016/j.ebiom.2023.104889
https://enlight-statement.org/

The official checklist and Explanation & Elaboration documents state a CC-BY-NC-ND licence. Original authorship, wording and licence notice remain on the source pages. The appended responses were supplied by the person completing this wizard.

Generated at https://resources.tscnlab.org/enlight/ . Completion indicates that checklist items have been addressed, not that a study has passed a quality assessment.
