## Supplementary figures and images for "Light-induced melatonin suppression in school-aged children before and during camping: associations with circadian phase and daytime light exposure"

### Supplemental Figure 1; Fig. S1

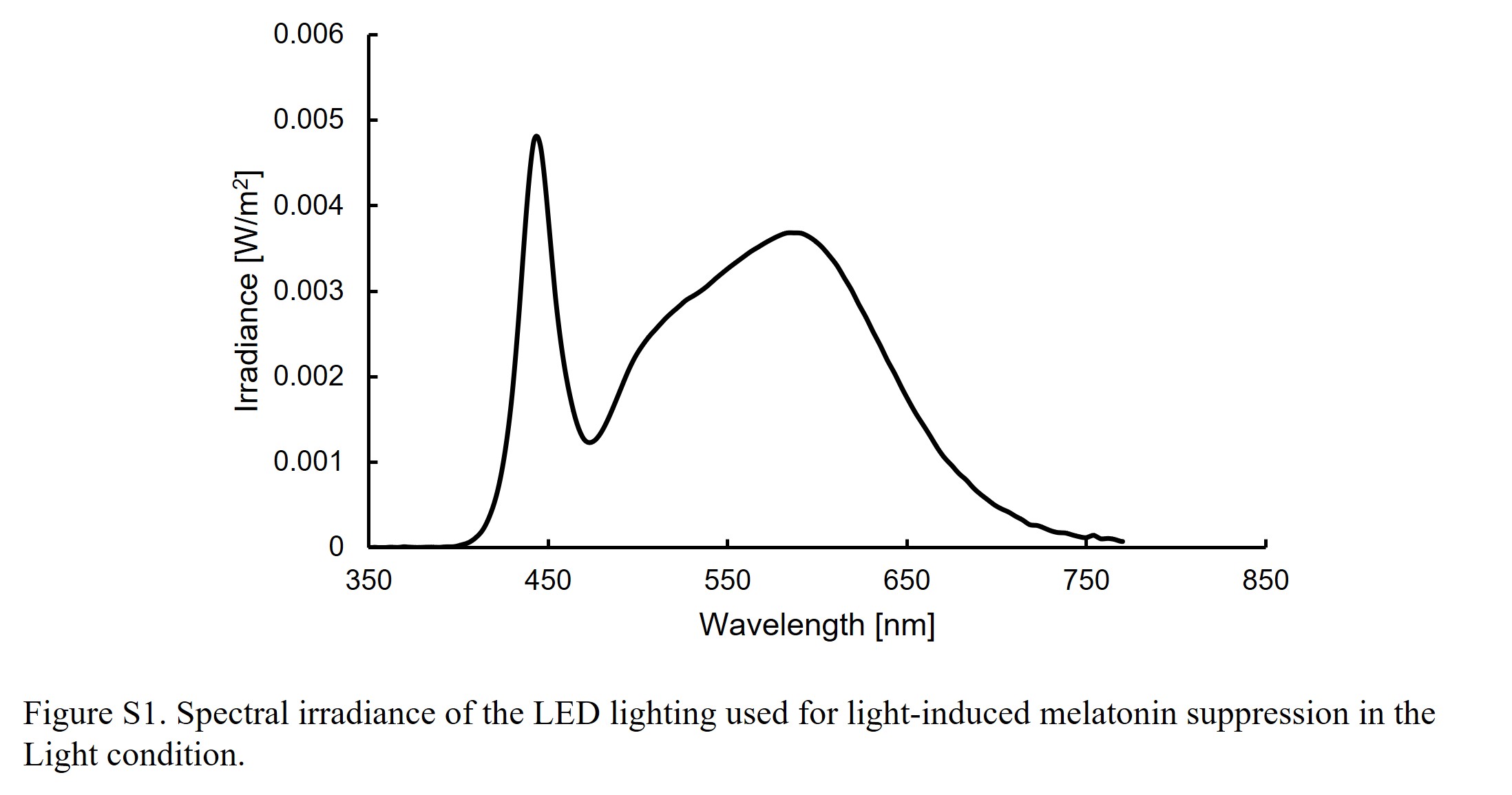

### Supplemental Figure 2; Fig. S2

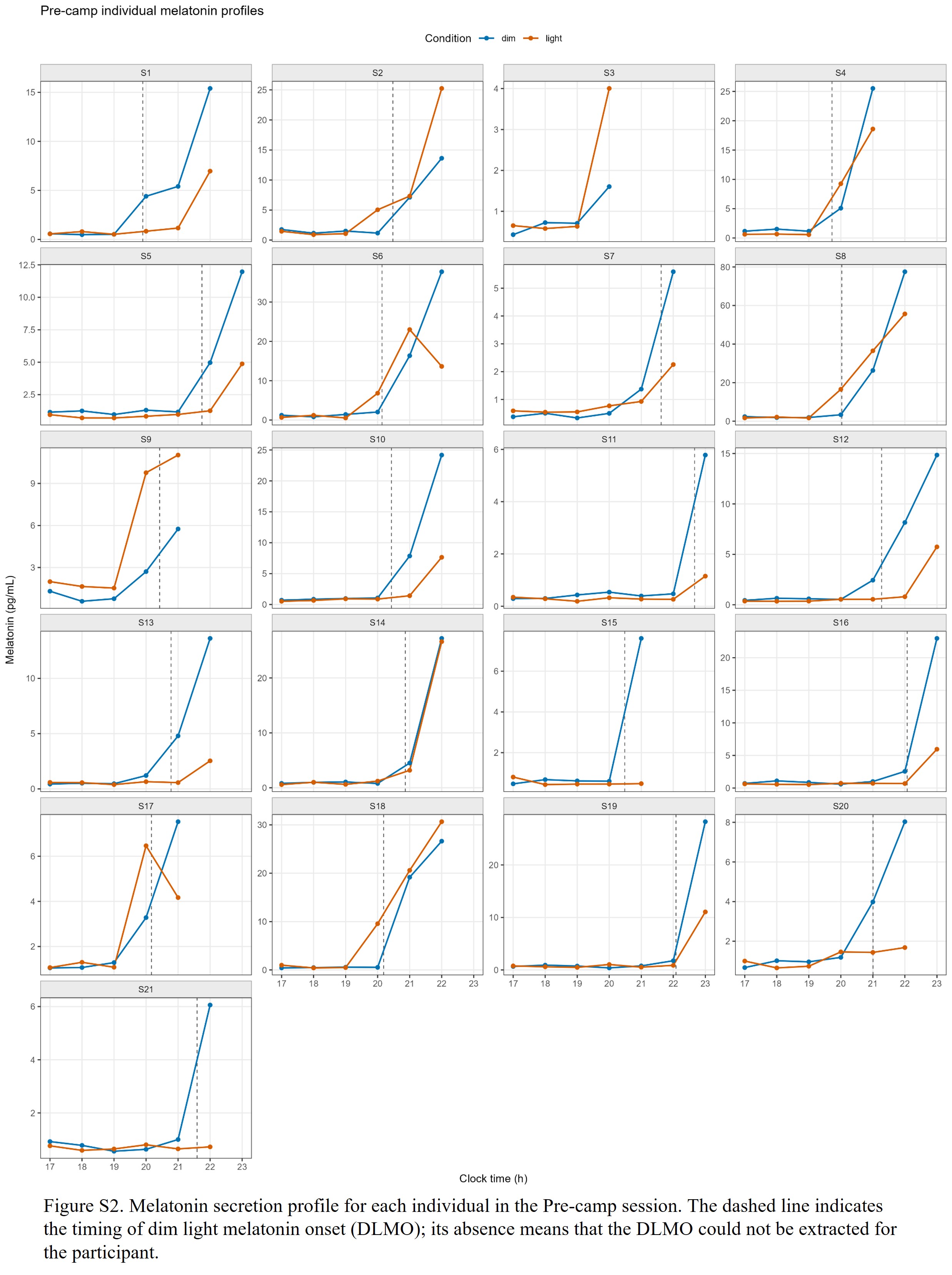

### Supplemental Figure 3; Fig. S3

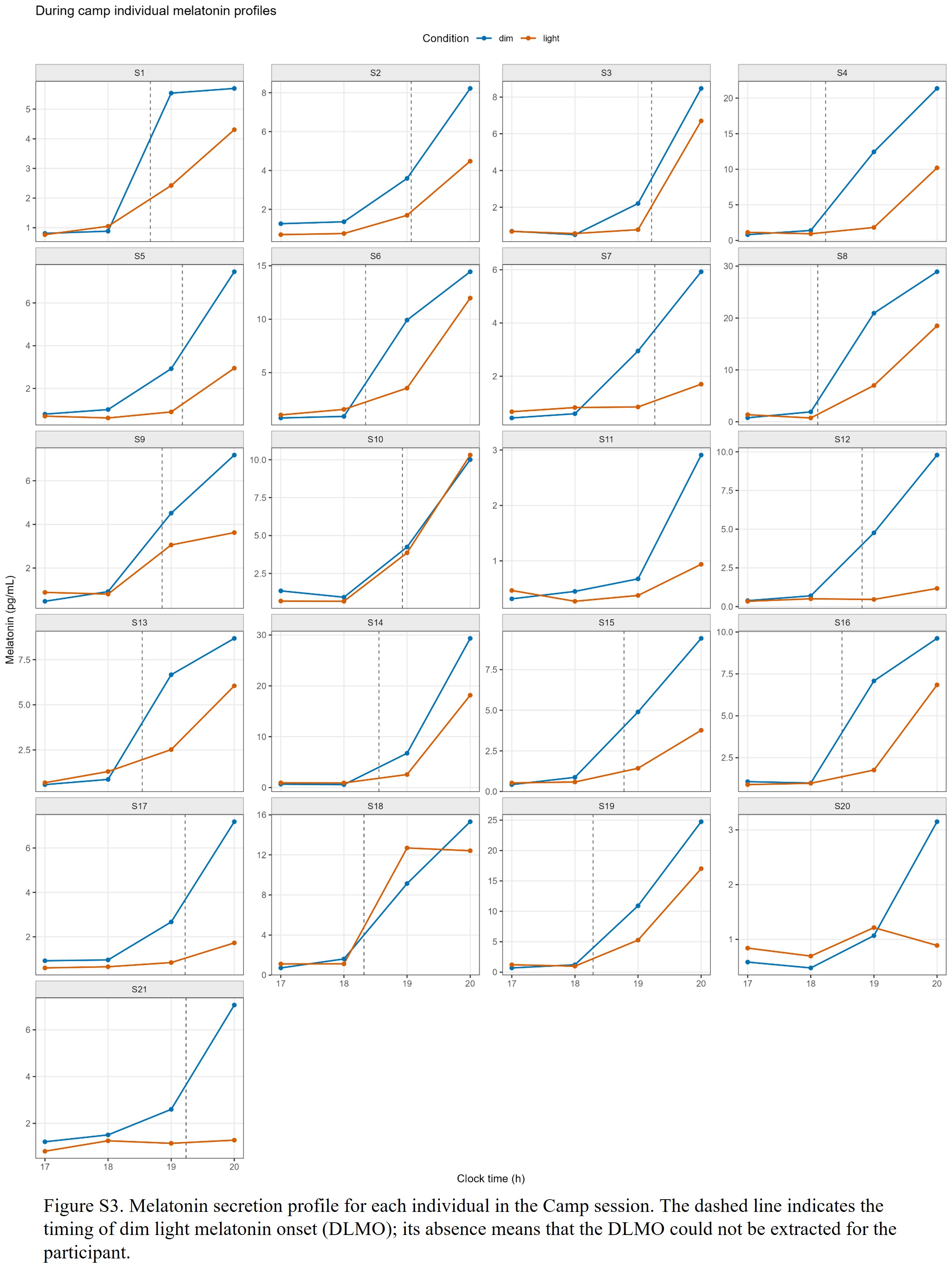

### Supplemental Figure 4; Fig. S4

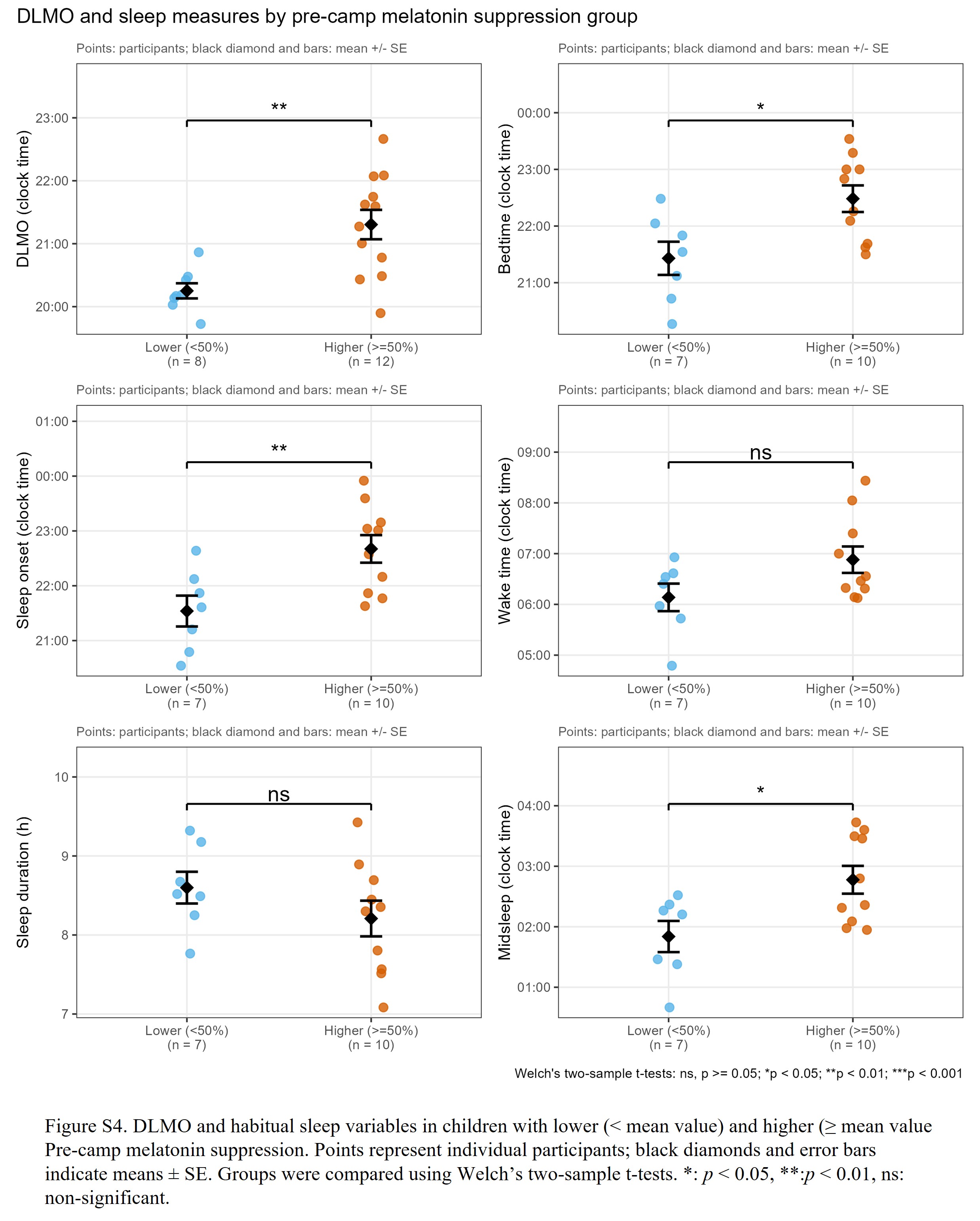
